# All-optical observation on activity-dependent nanoscale dynamics of myelinated axons

**DOI:** 10.1101/2022.07.18.500408

**Authors:** Junhwan Kwon, Sungho Lee, Yongjae Jo, Myunghwan Choi

## Abstract

In the mammalian brain, rapid conduction of neural information is supported by the myelin, whose functional efficacy shows steep dependence on its nanoscale cytoarchitecture. Although previous in vitro studies suggested that neural activity accompanies nanometer-scale cellular deformations, it has remained unexplored whether neural activity can dynamically remodel the myelinated axon due to the technical challenge in observing its nanostructural dynamics in living tissues. To this end, we introduced a novel all-optical approach combining a nanoscale dynamic readout based on spectral interferometry and optogenetic control of neural excitation on a living brain slice preparation. In response to optogenetically evoked neuronal burst firing, the myelinated axons exhibited progressive and reversible spectral redshifts, corresponding to the transient swelling at a subnanometer scale. We further revealed that the activity-dependent nanostructural dynamics was localized to the paranode. In summary, our novel all-optical studies substantiate that myelinated axon exhibits activity-dependent nanoscale swelling, which potentially serves to dynamically tune the transmission speed of neural information.

**RESEARCH SUMMARIES:** As neural activity involves rapid ion flux across the cell membrane, researchers have long been tried to detect the accompanying nanoscale morphological dynamics. However, measuring the activity-dependent nanostructural dynamics in the living mammalian brain has been an enigma due to the technical limitations. By combining excitatory optogenetics and *in situ* nanoscale metrology based on spectral interference, we demonstrate the first direct observation that the mammalian axons exhibit transient activity-dependent swelling at subnanometer-scale.

## INTRODUCTION

As neurons function by millisecond-scale ion flux across the cell membrane, neural activity has long been thought to accompany measurable morphological changes^1–3^. Since the late 1970s, several groups have reported nanometer-scale swelling in the giant axons of invertebrate species (crayfish and squid) by Michelson interferometry^4^ and mechanoelectrical measurement^5^. Later studies in mammalian cultured neurons by dark-field microscopy and full-field interferometry revealed the subnanometer-scale morphological dynamics dependent on electrical potential across the neuronal cell membrane^4–8^.

Yet, it is remained as an enigma to what extent neural activity dynamically remodel the axons in living mammalian brains, which are often ensheathed by the insulating layer - myelin. Myelin is a highly compacted subcellular structure of the oligodendrocyte, composed of multi-layered lipid membranes and intervening aqueous mediums. This highly-organized thin-film cytoarchitecture supports rapid and energy-efficient conduction of neural information in a small form factor, enabling formation of highly integrated circuit of the mammalian brains^9,10^. Considering that the function of myelin has steep dependence on its thin-film structure, the structural dynamics of myelinated axons, even at subnanometer-scale, can have critical impact on neural circuit functions^11^.

Investigating the nanostructural dynamics of myelinated axons in living mammalian brains is technically challenging. The current gold-standard on nanoscale imaging of myelinated axons is electron microscopy, which is hardly adoptable to the living biological samples^12–14^. Super-resolution techniques are a promising alternative for living biological samples but subnanometer-scale precision has yet been attained in the mammalian axons due to high optical aberrations of the lipid-rich myelin layers^15–17^. Thus, most studies so far focused on long-term dynamics of relatively large morphological changes involving cell proliferation and differentiation^11–13,18^.

Several years ago, we developed a spectral interferometric technique, named SpeRe, which offers the nanoscale readout of the thin-film cytoarchitecture of the myelinated axons in vivo^19^. Here, we combined the SpeRe’s nanoscale readout with optogenetic manipulation of neural activity to unveil the neural activity dependent nanostructural dynamics of myelinated axons in a living brain tissue. Our novel all-optical approach revealed that the myelinated axons exhibit subnanometer-scale swelling in response to neuronal burst firing, and that the swelling dynamics is cumulative and reversible in second-scale.

## RESULTS

### System for all-optical investigation

To observe morphological dynamics of functionally active myelinated axons at nanoscale, we introduced a novel all-optical neurophysiology approach (**Fig. 1a,b**). For manipulating neural excitation with minimal mechanical perturbation, we introduced red-shifted excitatory optogenetic protein (ChrimsonR) into cortical excitatory neurons. Neuronal excitation was timely triggered by epi-illuminated fiber-coupled LED at 633 nm on an acute brain slice, and was confirmed by GCaMP-mediated functional calcium imaging on the targeted neurons by two-photon microscopy^20^. For recording nanostructural dynamics of myelinated axons, we incorporated spectral reflectometry (SpeRe), that captures broadband reflectance spectrums at the geometric center of the axons and decodes the physical size of the multilayered thin-films (i.e., diameter of the myelinated axons) by decoding the spectrums. To assist the pinpointing of the geometric centers, we additionally introduced spectral confocal reflectance imaging (SCoRe)^21^, which provides a volumetric image of reflected light corresponding to the centerlines of myelinated axons.

**Figure 1.**
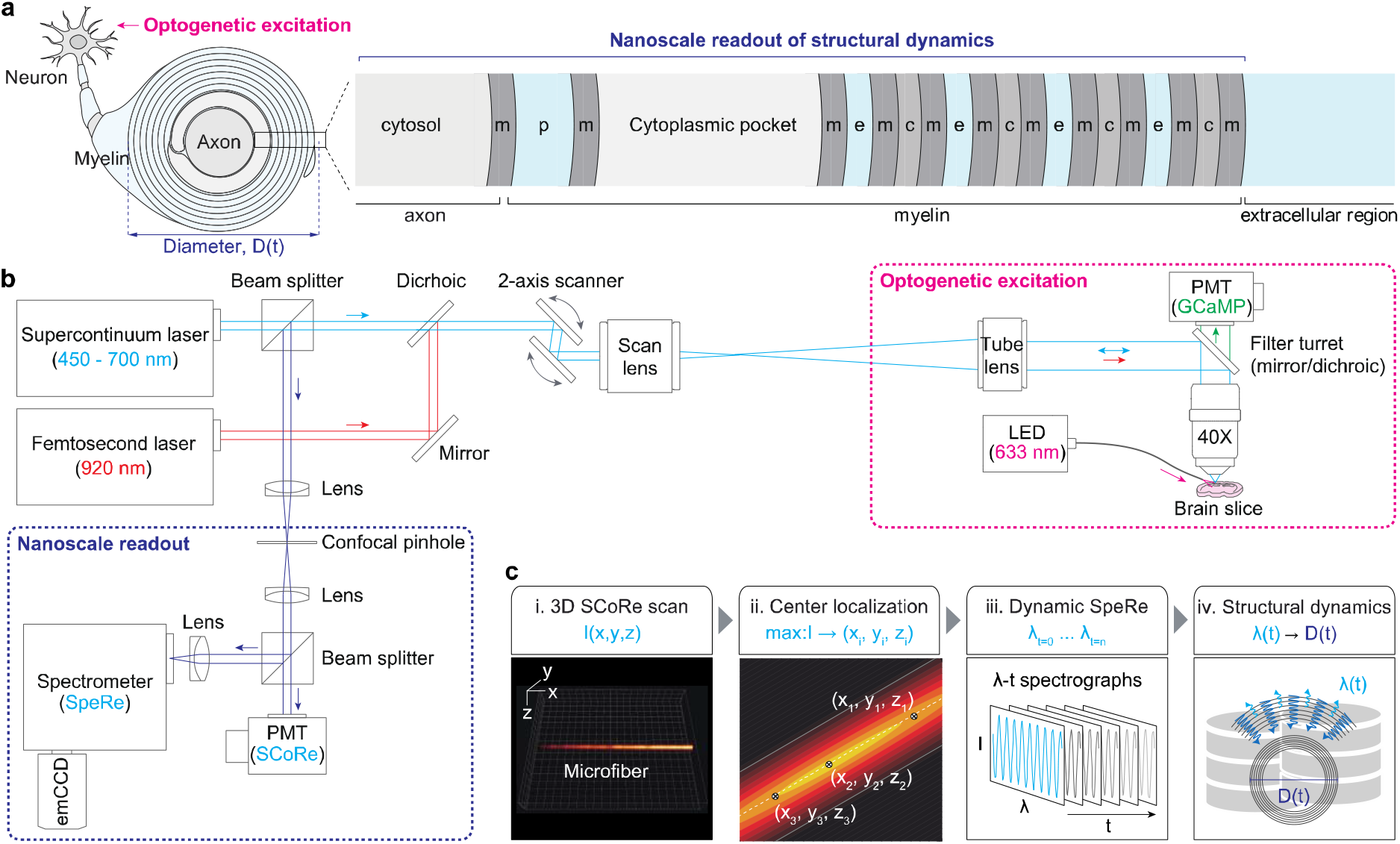
All-optical approach for observing activity-dependent nanostructural dynamics of living myelinated axons. (**a**) A cross-sectional view on the nanoscale cytoarchitecture of a myelinated axon at the paranodal region. The thin-film layers of the myelinated axons are shown in the magnified view. m, membrane. p, periaxonal space. c, cytosolic layer of the myelin. e, extracellular region. D, diameter. t, time. (**b**) The optic setup incorporating spectral confocal reflectance imaging (SCoRe), spectral reflectometry (SpeRe), two-photon fluorescence imaging, and optogenetic excitation. The supercontinuum laser serves as a light source for SpeRe and SCoRe, and the femtosecond laser provides two-photon excitation for recording GCaMP-mediated neuronal calcium activity. The auxiliary fiber-coupled light-emitting diode (LED) at 633 nm was used for triggering ChrimsonR-mediated excitatory optogenetics. (**c**) A pipeline for the nanoscale readout of structural dynamics of a myelinated axon. i, 3D SCoRe scan to acquire a volumetric reflectance image of myelinated axons. ii, Localization of the geometric centers based on the maximal reflectance intensity. iii, Time-lapse acquisition of the reflectance spectrographs at the geometric center. iv. Decoding of the structural dynamics from the acquired spectrographs. I, intensity. λ, wavelength.

The overall procedure for nanostructural readout begins with acquisition of a volumetric reflectance image from a fresh brain slice by SCoRe (**Fig. 1c**). From the volumetric SCoRe image, the position of maximum intensity for each cross-section is localized, resulting in the geometric centerline along the fibrous structure. The broadband reflectance spectrum is subsequently acquired over time at the center positions, and the structural dynamics is decoded from the acquired spectrums (SpeRe). For decoding of nanostructural dynamics, we performed numerical optic simulation on the myelinated axons and obtained quantitative relationship between the reflectance spectrum and the nanoscale cytoarchitecture. Briefly, interaction of light waves at the subcellular layers of myelinated axons were described by the thin-film matrix theory, and distribution of light waves at the focus was formulated by the vector diffraction theory. As the morphological dynamics was observed exclusively at the paranode of myelinated axons in our following experiments, we set the simulation parameters including physical size and refractive index for each subcellular layer based on the paranodal region (**Supplementary Table 1**).

The resulting simulation database showed that the swelling of myelinated axons (ΔD > 0) leads to negative phase-shift in wavenumber domain (Δv < 0) with linear relationship (**Supplementary Fig. 1**). Thus, we decided to use the relative phase-shift (Δv) as a reliable metric to quantify the dynamic swelling of myelinated axons (i.e., phase detection method^22–24^). To detect subnanometer-scale morphological dynamics, we used the grating of 600 lines·mm^-1^ and slit size of 10 μm, resulting in spectral resolution of ∼0.13 nm.

### Validation of subnanometer-scale readout precision

To verify subnanometer-scale precision of our SpeRe readout, we used a finely tapered glass fiber prepared by thermal drawing of a glass rod using a micropipette puller (**Fig. 2**). In a tapered fiber, the diameter (D) is a smooth function of its longitudinal position (x), therefore we can simply mimic the nanoscale change in diameter (ΔD) by shifting the position (Δx) using a galvanometric scanner. We first estimated the diameter along the fiber using an image acquired by polarization microscopy, and validated by SpeRe (**Fig. 2a-c**). We then selected two points, x_1_ and x_2_, which showed physiological axon diameters (0.3–5 μm). By the linear regression near the selected points, we acquired the slope (dD/dx), which provided the multiplicative factor for converting the step shift in position (Δx) to the change in diameter (ΔD). For example, to introduce increase in diameter by 0.5 nm at x_2_, we shifted the position along the centerline by +55 nm, corresponding to ΔD (0.5 nm) divided by dD/dx (0.009).

**Figure 2.**
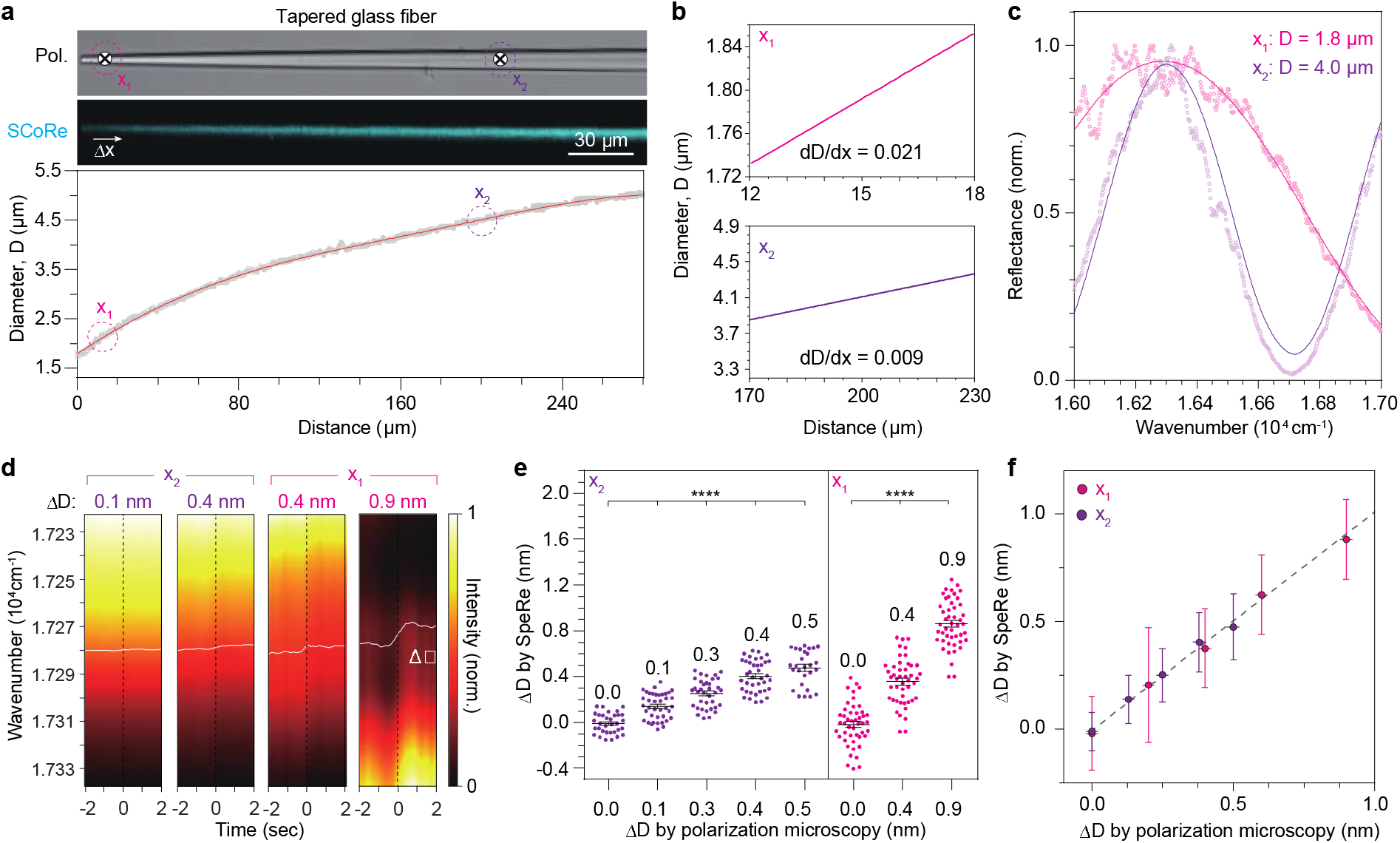
Validation on nanometer-scale readout precision of SpeRe. (**a**) A tapered glass fiber sample for quantifying the nanoscale readout precision. The tapered fiber is imaged by polarization microscopy (Pol.) and spectral confocal reflectance imaging (SCoRe). The gradually varying diameter (D) along the fiber was acquired from the polarization microscopic image. (**b**) The spatial derivatives of the diameter (dD/dx) at the representative positions (x_1_ and x_2_) along the tapered glass fiber. (**c**) Reflectance spectrographs at the positions, x_1_ and x_2_. The acquired spectrographs (dots) were overlaid with best-fit simulated spectrums (curved lines). (**d**) Representative time-lapse reflectance spectrographs measured at the indicated positions (x_1_ for ΔD = 0.4 nm and 0.9 nm; x_2_ for ΔD = 0.1 nm and 0.4 nm) with step shifts in position at 0 s. The step change in position was introduced by shifting the focal point along the centerline of the fiber. The white curves indicate the relative spectral shift. (**e-f**) Change in diameters (ΔD) estimated by SpeRe with the corresponding estimates by polarization microscopy. *, p < 0.0001 (unpaired t-test). The error bars indicate standard deviations.

In a tapered fiber, we obtained broadband reflectance spectra over time on the two selected positions at sampling speed of 20 Hz. To convert the spectral shift (Δv) to the change in physical diameter (ΔD), we applied the linear relationship derived from our numerical simulation for each position (**Supplementary Fig. 2a-c**). The in-position stability of SpeRe acquisition during the initial 2 s was approximately ±0.2 nm at x_1_ and ±0.1 nm at x_2_ in standard deviation (**Supplementary Fig. 2d-f**). The step shifts in the scan position (Δx) with variable distances were introduced at 2 s after the acquisition (**Fig. 2d-f**). By the step shift in the scan position, we observed the reliable negative spectral shift in wavenumber domain (Δv < 0), corresponding to the swelling. The estimated changes in diameter by SpeRe were precisely matched with the inputs introduced by the shift in position (**Fig. 2e-f**). These results suggest that our dynamic SpeRe readout provides subnanometer-scale precision.

### Activity-dependent nanostructural dynamics of myelinated axons

For optical manipulation and recording of neural excitation, we microinjected two types of adeno-associated viruses encoding a red-shifted opsin (AAV9-hSyn-ChrimsonR-tdTomato) and a fluorescent calcium indicator (AAV9-hSyn-GCaMP6s) at the somatosensory cortex of a live mice, and prepared fresh brain slices in oxygenized medium on the experimental day^25,26^ (**Fig. 3a**). By combining two-photon fluorescence and confocal reflectance imaging of the brain slice, we could visualize transfected neurons, as well as their myelinated axons. By illuminating 633 nm light pulses following the high frequency stimulation protocol, we observed reliable functional calcium activity in most neuronal soma expressing both ChrimsonR and GCaMP6s^10,13^ (**Fig. 3b** and **Supplementary Fig. 3**).

**Figure 3.**
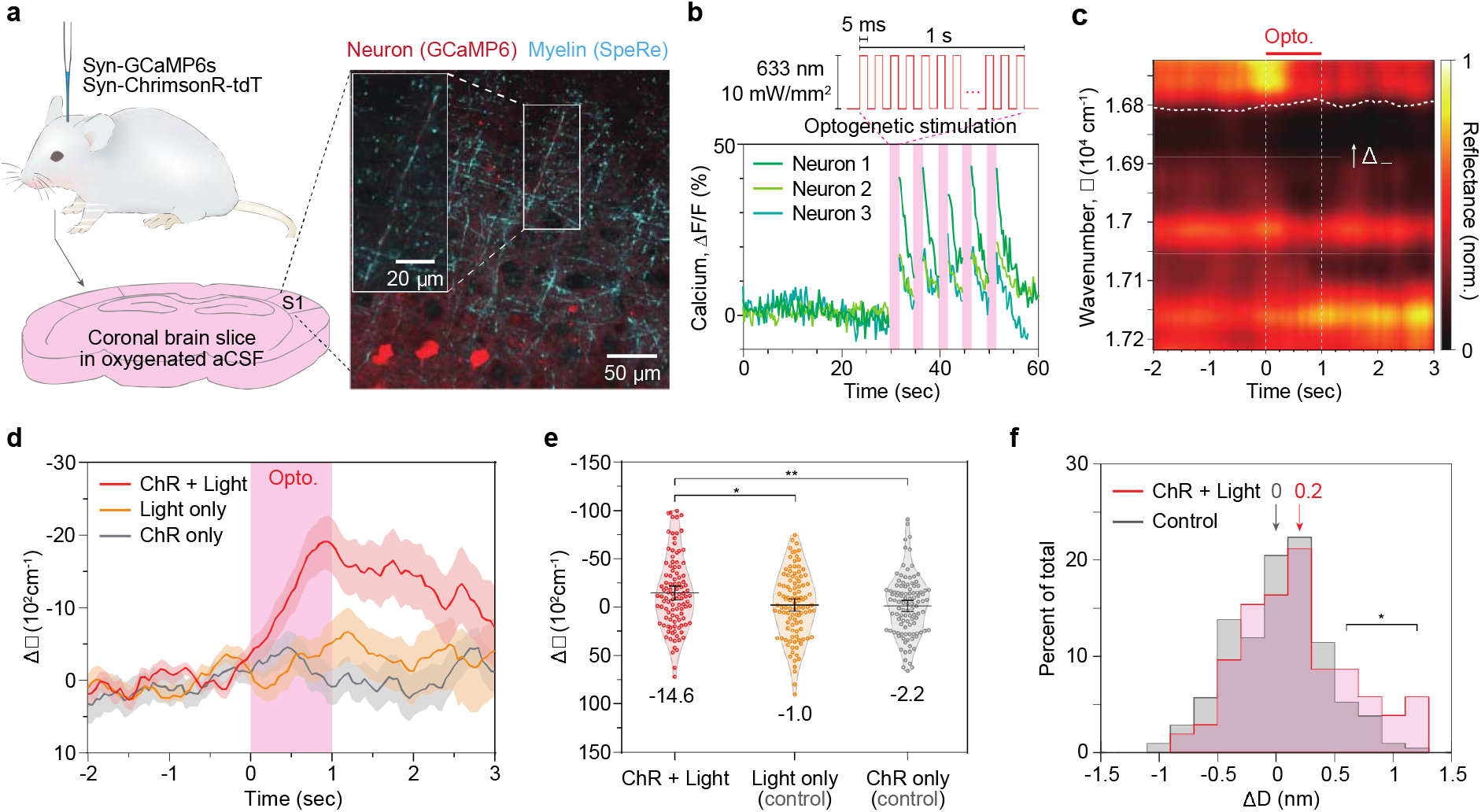
All-optical observation of activity-dependent subnanometer-scale swelling of myelinated axons. (**a**) Sample preparation. The neurons in the somatosensory cortex (S1) were transfected with a fluorescent calcium indicator (GCaMP6s) and an optogenetic actuator (ChrimsonR). A representative fluorescence/reflectance image from the brain slice is shown on the right. (**b**) Optogenetic excitation. Neurons transfected with ChrimsonR was excited by illuminating pulsed light (wavelength: 633 nm, irradiance: 10 mW/mm^2^, frequency: 100 Hz, pulse width: 5 ms). Calcium traces in soma from 3 neurons in (a) are shown. The shaded areas in pink indicate the duration of optogenetic stimulation. (**c**) A representative phase shifts (Δ□) induced by the optogenetic neural activity. The red bar indicates the duration of optogenetic stimulus (1 s). The white dashed line indicates the relative phase shift from the baseline (-1.0 – -0.5 s). (**d**) The group averaged phase shifts (Δ□) for each experimental group. The red curve (‘ChR + Light’ group, n = 104 axons in 5 mice) indicates the ChrimsonR-expressing neurons with light stimulus, the yellow curve (‘Light only’ group, n = 105 axons in 5 mice) indicates the wild-type neurons receiving photostimulation, and the gray curve (‘ChR only’ group, n = 105 axons in 5 mice) indicates the ChrimsonR-expressing neurons without light stimulus. Note that only the ‘ChR + Light’ group exhibited distinguishable negative shift in wavenumber. (**e**) Statistical group comparison of phase-shifts (Δ□) in (d). The phase-shifts were quantified by averaging the relative phase shifts during 0.5–1 s. *, p < 0.05 (unpaired t-test). **, p < 0.01 (unpaired t-test). (**f**) Histograms of activity-dependent changes in diameters of myelinated axons (ΔD). The positive value of ΔD corresponds to swelling.

To address whether neuronal excitation leads to nanostructural dynamics of myelinated axons, we randomly sampled up to 10 myelinated axons for each brain slice, and recorded the nanoscale dynamics by SpeRe for 5 s with light stimuli (‘ChR + Light’ group, n = 104 axons in 5 mice; **Fig. 3c,d**). As negative control groups, we included brain slices without light stimuli (‘ChR only’ group, n = 105 axons in 5 mice) and also brain slices without introducing the optogenetic actuator (‘Light only’ group, n = 105 axons in 5 mice). The SpeRe readout was performed at near the either ends of myelin sheaths, corresponding to the paranode, where axo-myelinic communication is known to be active^27–29^. Apparently, only the group with functional optogenetic excitation (‘ChR + Light’ group) showed statistically significant spectral shift by ∼2,000 cm^-1^ on average (unpaired t-test: p < 0.05; **Fig. 3e**). Intriguingly, the spectral shift was cumulatively increased during the 1 s period of optogenetic excitation and slowly recovered in several seconds, indicating that the activity-dependent morphological dynamics does not follow the neuronal membrane potential having millisecond-scale rise and fall kinetics. Moreover, pharmacological inhibition of action potential generation by tetrodotoxin significantly, but only partially, attenuated the swelling, suggesting that the generation of action potentials is not necessary for inducing the swelling (**Supplementary Fig. 4**).

**Figure 4.**
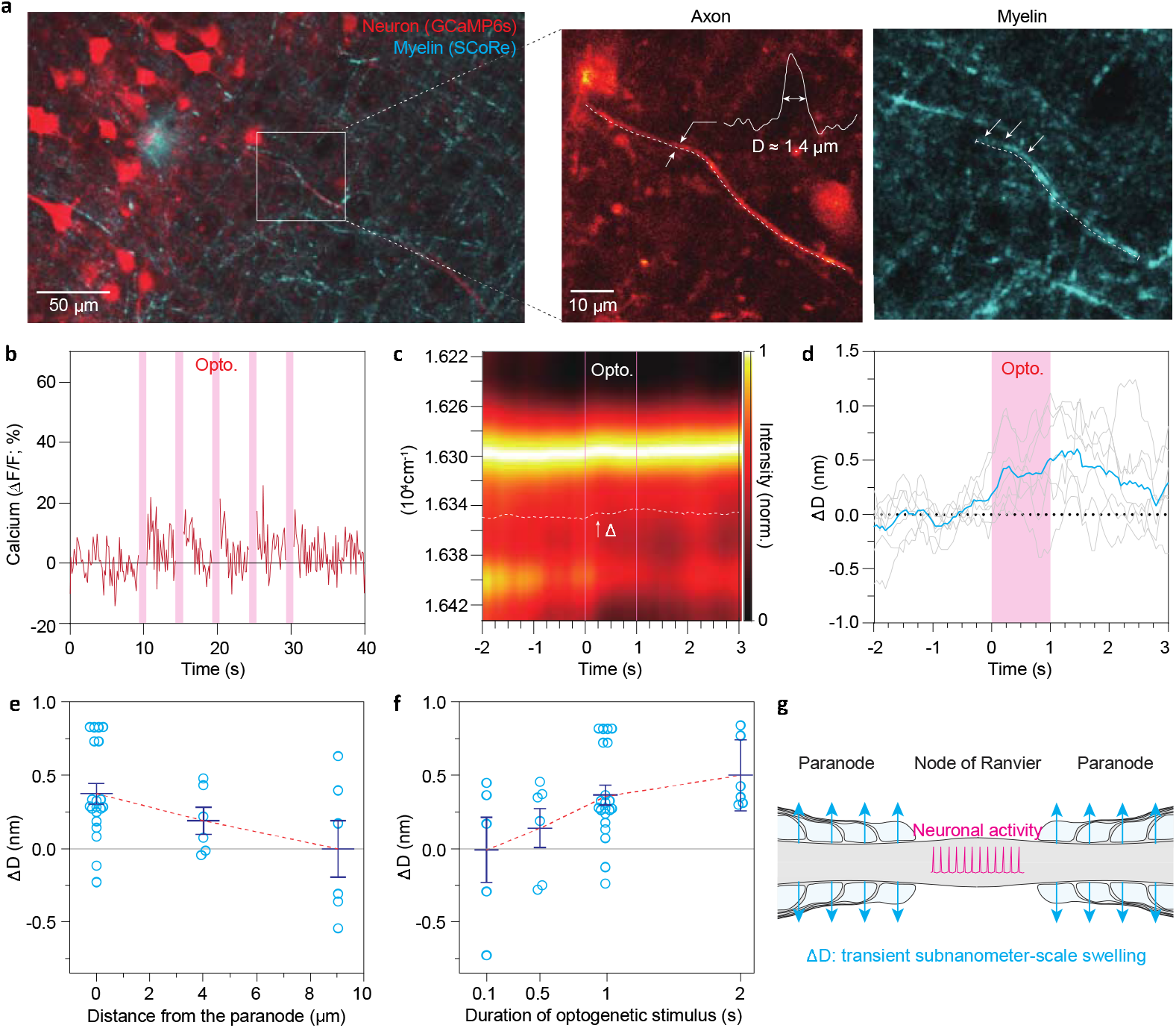
Correlative characterization of subnanometer-scale swelling in an optogenetically active axon. **(a)** Representative images of myelinated axons. The neurons are imaged by two-photon fluorescence of GCaMP6s (red) and the myelin is visualized by SCoRe (cyan). The magnified view of the myelinated axon of interest is shown on the left. The axon diameter (D) was estimated to by full-width-half-maximum of the transverse intensity profile (D ≈ 1.4 μm). The dashed lines indicate the axon and the myelin sheath. **(b)** Axonal calcium activity triggered by optogenetic excitation of ChrimsonR. The shaded area in magenta indicates the duration of optogenetic excitation (633 nm, 10 mW/mm^2^). **(c)** Observation of time-dependent change in reflectance spectrum at the optogenetically active myelinated axon. The dashed lines indicate the duration of optogenetic stimulation. Note the progressive spectral redshift (Δ□) during the optogenetic stimulation. **(d)** Time dependent changes in the diameter of myelinated axons (ΔD) in response to optogenetic stimulation (shaded region in magenta). Cyan curve indicates the averaged trace of 7 repeated trials (individual traces are shown in grey). **(e)** The swelling is localized to the paranode. **(f)** Dependency of ΔD on the duration of optogenetic stimuli at the paranode. (**g**) The schematic diagram of the activity-dependent nanoscale dynamics of the myelinated axons.

To convert the spectral shift (Δv) to the change in physical diameter (ΔD), we applied the linear relationship derived from our numerical simulation (**Supplementary Fig. 1**). As the nanostructural parameters for individual myelinated axons we sampled were not attainable, we applied the representative structural parameters in **Supplementary Table 1** obtained from previous electron micrographs and estimated from our optical images (**Supplementary Fig. 3**)^19,30^. Although this approach compromised the precision of estimation on individual axons, we were able to gain information on the population distribution.

Although statistically significant, the group-averaged spectral shift of ∼2,000 cm^-1^ corresponds to ∼0.2 nm, which was at least several folds smaller than previously reported axonal swelling observed in cell culture and invertebrate systems (**Fig. 3e**). This discrepancy can be explained by our experimental design, which randomly samples myelinated axons in brain slices. Conceivably, a significant portion of long-projecting myelinated axons is expected to be severed during the tissue slicing procedure and contains insufficient optogenetic proteins due to stochastic nature of viral transfection. Consequently, the large portion of the samples even in the ‘ChR + Light’ group might have been nonresponsive to optogenetic stimuli. Indeed, the histograms of change in diameters for the experimental and the negative control groups were largely similar with statistically significant difference observed only at the swelling greater than ∼0.5 nm (**Fig. 3f**).

We thus questioned if we could observe reliable nanostructural dynamics in the myelinated axons that are morphologically intact and functionally active (**Fig. 4**). By two-photon imaging of neuronal morphology and functional calcium activity with optogenetic stimuli, we identified the three morphologically intact and functionally active myelinated axons, which were connected to the intact neuronal soma (**Fig. 4a,b**). As expected, the three myelinated axons repeatedly exhibited progressive increase in their diameters by 0.3–1 nm in response to optogenetic stimuli (**Fig. 4c,d** and **Supplementary Fig. 5**). We further observed that the morphological dynamics was localized to the paranode (**Fig. 4e**) and that the degree of swelling was positively correlated with the duration of optogenetic stimuli (**Fig. 4f**).

## DISCUSSION

By combining nanoscale readout based on spectral interferometry (SpeRe) and optogenetic manipulation of neural excitation, we reported the first experimental evidence that myelinated axons exhibit activity-dependent subnanometer-scale morphological dynamics of ∼0.5 nm. We further revealed that the nanostructural change displays slow kinetics at second-scale and localized to the paranode. As conduction efficacy of neural information has steep dependence on the subcellular structure of the paranode, we expect that its nanostructural dynamics can serve as a regulatory mode of controlling conduction speed in the mammalian brains.

The functional consequence of the morphological remodeling at the paranode can be inferred by the periaxonal nanocircuit model of the myelinated axons^10^. According to this model, the periaxonal space is electrically conductive and the paranodes are only partially sealed, resulting in leaky propagation of neural information. Thus, the swelling of the paranode can attenuate the leakage and accelerate the conduction speed. By the theoretical double cable model, 1 nm swelling at the paranode leads to ∼1.3% acceleration in conduction speed. Accordingly, the observed swelling of ∼0.5 nm is expected to increase the conduction speed by ∼0.7%. Consistently, Yamazaki et al. reported short-term increase in conduction speed by directly depolarizing the myelinating oligodendrocyte^28,31^. The physiological relevance of this change in conduction velocity on neural circuit function requires further investigation.

Our results revealed that the paranodal swelling occurs cumulatively and reversibly at second scale, suggesting that the neuronal membrane potential having millisecond-scale rise and fall kinetics is not the direct source of the swelling (**Fig. 3d** and **Fig. 4d**). In addition, pharmacologic inhibition of voltage-gated sodium channel by tetrodotoxin only partially attenuated the paranodal swelling (**Supplementary Fig. 4**), suggesting that generation of action potential is not necessary for inducing the swelling. Considering that ChrimsonR is light-gated channel permeable to both sodium and calcium ions^26,32^, we expect the calcium influx may play a key role in the paranodal swelling. In agreement with our thought, we observed calcium transients by ChrinsonR-mediated optogenetics in our inhibition study with tetrodotoxin, and the degree of calcium increase was attenuated compared to the control group. Further pharmacological inhibition of ion channel subtypes or use of ion-selective opsins will clarify the underlying molecular mechanism.

Although our results consistently suggest that the neuronal excitation leads to enlargement of myelinated axons at the paranode, it remains to be answered which subcellular components are remodeled. Chéreau et al. reported that high frequency stimulation of neurons leads to axonal swelling in the mouse brain^15^. Trigo and Smith also showed that micron-scale axonal swelling following prolonged electric stimulation of the peripheral nerves^27^. As mature myelin sheaths do not typically exhibit axonal activity dependent functional responses, we estimate that the axon at the node of Ranvier swells in response to ionic redistribution across the membrane and the surrounding myelin is passively enlarged. Detailed multiparametric analysis based on the thin-film model with a priori structural information on each subcellular composition may provide a solution to this question.

## METHODS

### Tapered glass fiber sample

The tapered glass fibers were obtained by the thermal drawing of a glass rod (1.5 mm in diameter) using a micropipette puller (P-1000, Sutter Instrument). To mimic the typical axon diameters (1–5 μm), the optimal parameters for the micropipette puller were obtained by trials and errors (ramp = 500 temperature = 550, pull = 40, velocity = 20, and pressure = 500). The tapered fiber was glued on a plain slide glass using an acrylic adhesive (401, Loctite) ensuring that the fiber is parallel to the slide glass (i.e., orthogonal to the optical axis of the objective lens). To firmly hold the fiber, a silicone-based sealant (Kwik-Cast, World Precision Instruments) was introduced around the fiber tip, exposing only the fiber tip in the air. The sample was imaged by a polarization microscope (SP8, Leica) and the diameter along the fiber was measured by applying the ‘Distance map’ module in ImageJ.

### Artificial cerebrospinal fluid (aCSF)

Two types of artificial cerebrospinal fluid (aCSF) solutions were prepared, one is for surgery and the other is for recording. The surgical aCSF was composed of (in mM) 92 NMDG, 2.5 KCl, 1.25 NaH_2_PO_4_, 30 NaHCO_3_, 20 HEPES, 25 glucose, 2 thiourea, 5 Na-ascorbate, 3 Na-pyruvate, 0.5 CaCl_2_·2H_2_O, and 10 MgSO_4_·7H_2_O. The recording aCSF was composed of (in mM) 124 NaCl, 2.5 KCl, 1.2 NaH_2_PO_4_, 24 NaHCO_3_, 12.5 glucose, 2 CaCl_2_·2H_2_O, and 2 MgSO_4_·7H_2_O (Sigma Aldrich). Both solutions were titrated to have pH 7.3–7.4 and 300–310 mOsm while aerated with 95% O_2_ and 5% CO_2_.

### Mouse preparation

All mice were housed with littermates in groups of two to five in reverse day/night cycle and given ad libitum access to food and water. All animal experiments were performed in compliance with institutional guidelines and approved by the sub-committee on research animal care at Sungkyunkwan University and Seoul National University. Male or female C57BL6J wild-type mice aged 4-week-old (Jackson Laboratory) were used for virus-mediated transduction of a genetically-encoded calcium indicator (GCaMP6s) and/or an optogenetic protein (ChrimsonR-tdT). The mouse was anesthetized by inhaling 4% isoflurane (Hanapharm) in an induction chamber and were subsequently maintained with 1– 1.5% isoflurane during surgery. The mouse skull was affixed on a custom-made stereotaxic frame and the body temperature was maintained at 37°C using a homeothermic blanket (TC-1000, CWE) and a thermistor probe (YSI-451, CWE). After removing the scalp, the hole was made on the center of the somatosensory area at a diameter of ∼1 mm. The 700 nL of a solution containing AAV9-hSyn-GCaMP6s and/or AAV5-hSyn-Chrimson-tdTomato (∼5 × 10^11^ GC·ml^-1^ each in the recording aCSF) was slowly infused to the cortical layer 3–4. After 3 weeks, the mice were used for the acute slice experiments.

### Brain slice preparation

Mice were decapitated under deep anesthesia by inhaling 3% isoflurane in O_2_. The mouse brain was harvested and sliced using a vibratome (thickness = 300 µm; VT1200S, Leica). During the slicing procedure, the immersion solution was the surgical aCSF solution kept at 4°C. Subsequently, the brain slices were incubated in the surgical aCSF solution at 35°C for 20 min and were immersed in the recording aCSF solution at room temperature (22–24°C) with continuous aeration of 95% O_2_ and 5% CO_2_ for 30min. The brain slices were mounted on an imaging chamber using a tissue anchor (SHD-41/10, Warner instruments). For the neuronal inhibition study, tetrodotoxin (TTX) was added to the recording aCSF at 10 μM.

### Optic setup

Our customized optic system shown in Fig. 1b was designed to incorporate the following 3 modalities: (i) two-photon fluorescence imaging for recording neuronal activity; (ii) spectral reflectance spectroscopy for nanostructural readout of myelinated axons; and (iii) optogenetics for manipulating neural activity. The system was constructed based on an upright galvanometer-based laser scanning microscope (Ultima IntraVital, Bruker), coupled to a Ti-Sapphire femtosecond laser (for two-photon fluorescence imaging; Chameleon Ultra II, Coherent) and a supercontinuum white-light laser (for SpeRe and SCoRe; EXB-6, NKT photonics). The femtosecond laser was tuned to 920 nm for exciting GCaMP6s and was attenuated to 10–20 mW at the objective back aperture. The supercontinuum laser was attenuated to ∼0.4 mW at the objective back aperture using a neutral density filter and bandpass-filtered to 450–700 nm. An apochromatic water-immersion objective lens (25X, 0.95 NA, Leica) was used for both two-photon fluorescence and SpeRe/SCoRe readouts. For two-photon fluorescence imaging, a GaAsP photomultiplier tube placed at the non-descanned path was used along with a bandpass filter at 500–550 nm. For the SCoRe imaging, a silicon photomultiplier tube placed at the descanned path was used. For SpeRe measurements, an array spectrometer (SR303i and Newton, Andor) was introduced at the descanned path. For spectroscopy, the grating of 600 lines·mm^-1^ was adjusted to accept spectral window of 550 to 650 nm, where the input white light exhibited near uniform intensity profile over the spectral window. The slit size was set to 10 μm, providing a spectral resolution ∼0.13 nm with enough signal-to-noise ratio at acquisition speed of 20 Hz. For optogenetics, a 633 nm diode laser (MRL-III-633, CNI laser) was coupled to a multimode fiber (400 µm core, 0.39 NA; M119L02, Thorlabs), which was mounted on a motorized 3-axis micromanipulator (MP-285, Sutter Instrument). Optical irradiance for optogenetic stimulation was set to 10 mW·mm^-2^ at the tissue surface, which was delivered at 100 Hz with 50% duty cycle for the duration of 0.5–2 s.

### Data analysis

For SpeRe, the time-series reflectance spectrums were filtered in spectral and time domains in Matlab. To reduce artifactual jittering noise, the lowpass filter at a cutoff frequency of 0.65 nm^-1^ was applied in the wavelength domain by applying the ‘lowpass’ function, and the smoothing filter with the bin of 0.5 s was applied in temporal domain by using the ‘smooth data’ function. From the filtered spectral data, relative phase shift over time from the baseline was retrieved based on the least-square method. Occasionally, unpredictable motion artifacts (e.g., instability of media perfusion) interfered reliable quantification of the phase shift. Thus, we excluded the data if it displayed at least one of the following indications of excessive motion artifact: change in reflectance intensity greater than 10% and the phase drifted over 3 nm. For calcium imaging data, we quantified relative change in fluorescence normalized by the baseline fluorescent intensity (ΔF/F).

### Statistical analysis

GraphPad Prism was used for statistical analysis. Group comparisons were conduction using unpaired t-tests (parametric). The data are presented as mean ± standard error. We considered a p-value less than 0.05 to be statistically significant.

### Code availability

The MATLAB scripts used for data analysis are available at an open source repository (https://github.com/Neurophoton).

### Data availability

All relevant data are available from the corresponding author upon request. A reporting summary for this Article is available as a Supplementary Information file.

## Acknowledgments

We thank Dr. Moonseok Kim for guidance on the optic simulation and Dr. Seong-gi Kim for the support on the experimental infrastructure. This work was supported by the Institute of Basic Science (IBS-R015-D1) and by the by Basic Science Research Program through the National Research Foundation of Korea (NRF) funded by the Ministry of Education (2018R1D1A1B07042834, 2019M3E5D2A01058329, 2019M3A9E2061789, 2020M3C1B8016137, 2020R1A5A1018081).

## Author contributions

M.C. initiated and supervised the study. Y.J. developed the optical hardware and acquisition software. J.K. performed the brain slice experiments and data analysis. All the authors cowrote the manuscript.

## Corresponding author

Correspondence to M. Choi

