## Supplementary material for "All-optical observation on activity-dependent nanoscale dynamics of myelinated axons": Supplemental files.docx

**Supplementary Figure 1**

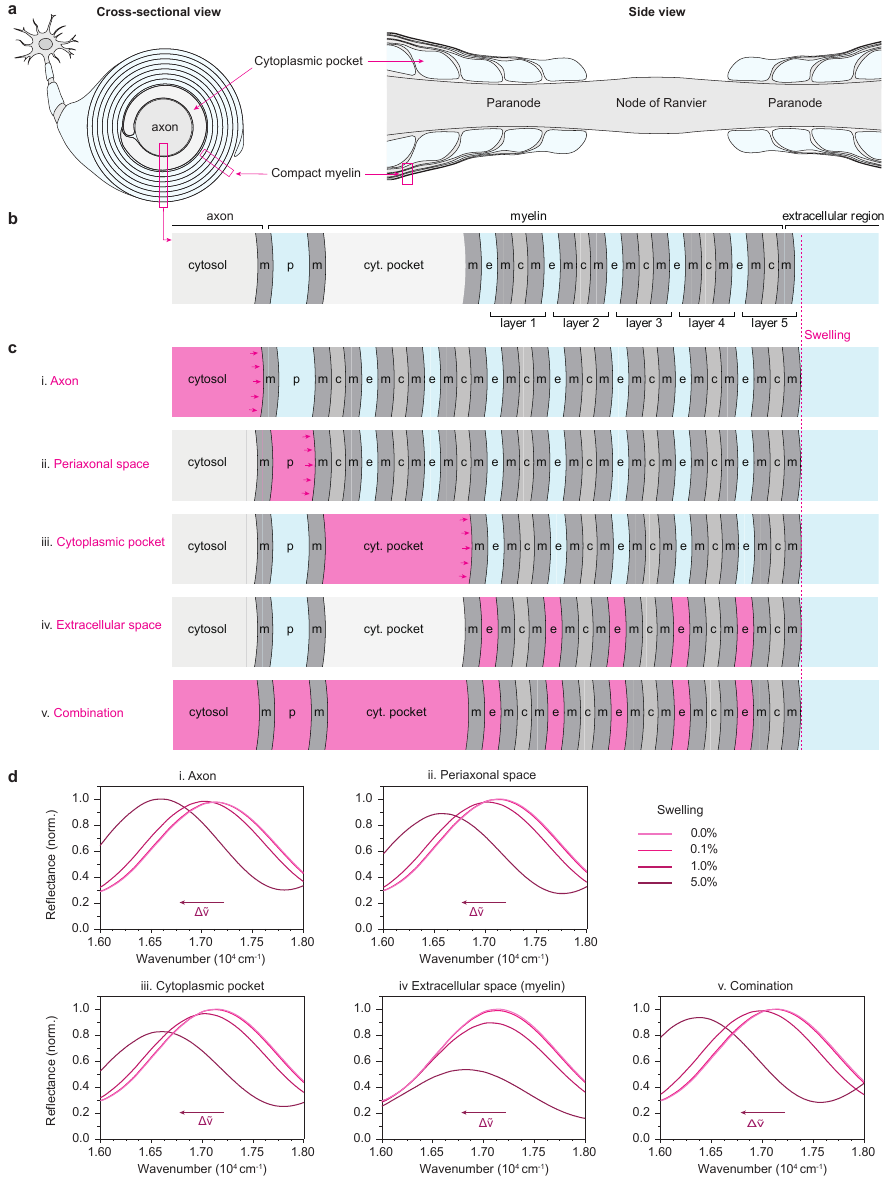

**Supplementary Figure 1 | Spectral shift represents nanoscale structural change.** (**a**) Schematic illustrations of a neuron with a myelinated axon fiber. Left, a cross-sectional view of a myelinated axon at the paranodal region. Right, a side view of a myelinated axon at the node and paranodal regions. Note that the cytoplasmic pocket is enlarged at the paranodal region. (**b**) A thin-film model for the paranodal region. (**c**) Potential swelling modes of myelinated axons. The affected subcellular regions are shown in magenta. i, the axon is swollen. ii, the periaxonal space is enlarged. iii, the cytosolic pockets are swollen. iv. the extracellular spaces of myelin is swollen. v. combination of the above. (**d**) Spectral shift in SpeRe for the 5 cases shown in (c). Note that for all the cases, the swelling in the cavity size leads to the spectral redshifts in the similar degree, suggesting that the spectral shift can serve as a reliable indicator for tracing the nanoscale swelling dynamics of myelinated axons.

**Supplementary Figure 2**

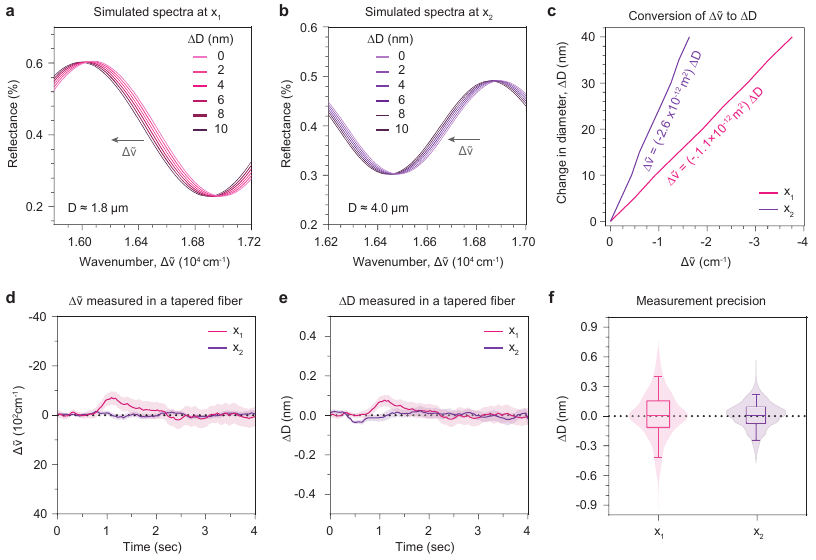

**Supplementary Figure 2 | Analysis pipeline for the stable SpeRe acquisition.** (**a,b**) In a tapered glass fiber sample shown in Fig. 2a, numerical simulation produced reflectance spectrums while increasing the diameters with 2 nm increments for the two locations (base x_1_, 1.8 μm and x_2_, 4 μm). (**c**) the linear relationships between the spectral shifts (Δṽ) and the diameter increases (ΔD) acquired from (**a,b**). (**d-f**) the SpeRe acquisition was performed at a sample rate of 20 Hz for a period of 6 s. (**d**) Artifacts in spectral shift over time. The SpeRe acquisition (Δṽ, relative shift in wavenumber) was performed at the two locations on the tapered fiber indicated in Fig. 2a (x_1_ and x_2_). (**e**) Converting the spectral shift (Δṽ) to the change in diameter (ΔD). Using simulation database, the relationship between Δṽ and ΔD was deduced and applied. (**f**) Precision of the dynamic SpeRe acquisition. The stability in standard deviation at x_1_ was 0.2 nm and that at x_2_ was 0.1 nm.

**Supplementary Figure 3**

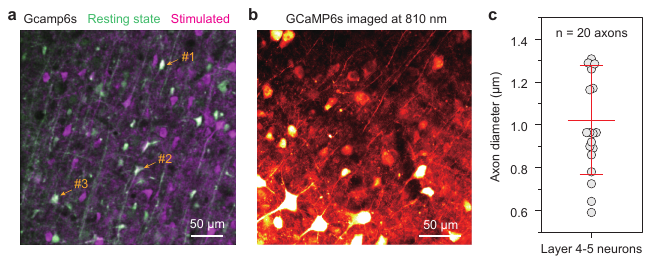

**Supplementary Figure 3 | Physical parameters of the myelinated axons.** (**a**) A representative image of a mouse brain slice virally transfected with a genetically encoded calcium indicator and an optogenetic actuator. Resting state neurons (green) and activated neurons by optogenetic light (magenta) of GCaMP6s imaged by two-photon at the functional imaging wavelength (920 nm). (**b**) Anatomical image of neurons and their axon fibers. This pseudo-color image is acquired from two-photon fluorescence of GCaMP6s excited at the isosbestic wavelength (810 nm). (**c**) Quantified axon diameters from the image acquired in (**b**). Axon diameters originated from the layer 4-5 neurons in the somatosensory area were acquired by quantifying full-width-half-maximum of the transverse intensity profile (n = 20 axons). The error bar indicates standard deviation.

**Supplementary Figure 4**

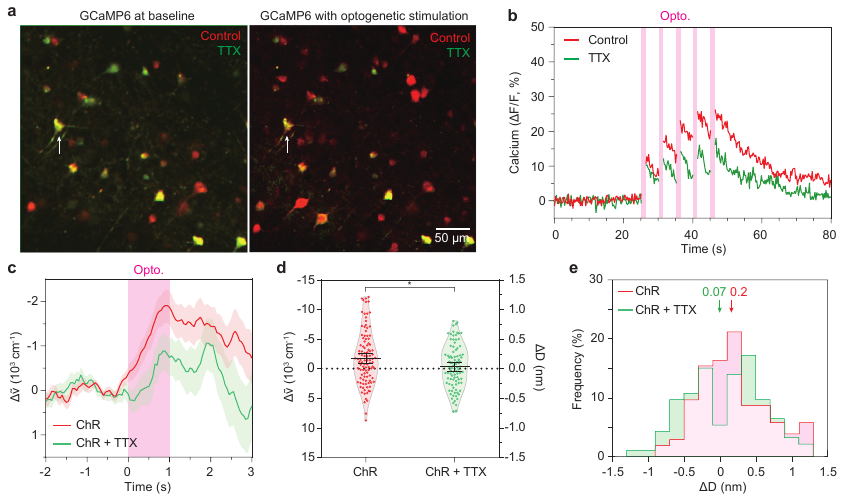

**Supplementary Figure 4 | Inhibition of voltage-gated sodium channel by tetrodotoxin (TTX) only partially inhibited the activity-dependent swelling of myelinated axons.** A brain slice from a virally transfected with GCaMP6s and ChrimsonR-tdTomato was bathed with 0 or 10 μM TTX in aCSF. (**a**) Merged Gcamp6s image of neurons taken by two-photon imaging. Left, Resting state of Gcamp6s signal. Right, Activated neurons by optogenetic stimulation. Red, Neurons in aCSF with no TTX. Green, Neurons in aCSF with 10 μM TTX. Both images exhibited attenuated Gcamp6s signals by TTX. (**b**) T-series calcium changes. Calcium transient of neurons in TTX medium exhibited attenuated amplitude in ΔF/F. (**c**) Comparison of time-series phase shift. Pink shadow, period of optogenetic stimulation. Solid line, averaged trace for each group. Shadow, standard error of mean. (**d**) Maximum phase difference between mean of baseline (-1 to -0.5) and mean of stimulated period (0.5 to 1) in (c). (**e**) Comparison of distributions of maximum diameter changes estimated by SpeRe. TTX, n = 93 myelinated axons from 4 mice.

**Supplementary Figure 5**

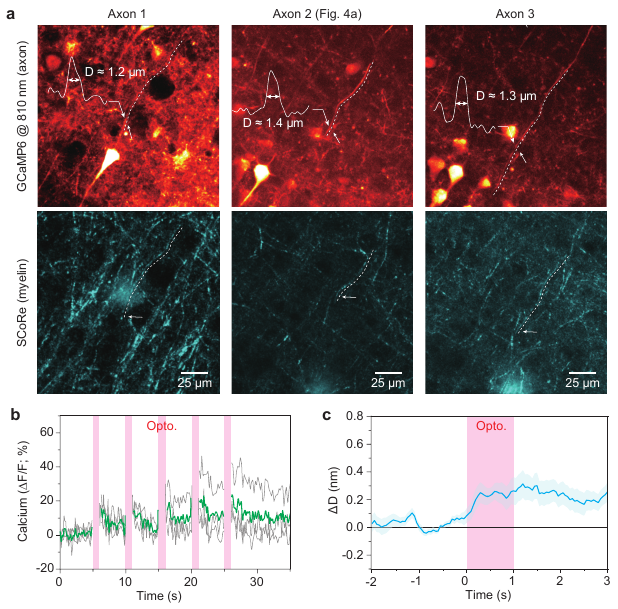

**Supplementary Figure 5 | Correlative observation on activity-dependent subnanometer-scale swelling of optogenetically active myelinated axons.** (**a**) The representative cases of optogenetically active myelinated axons (n = 3 axons). Upper, axons imaged by two-photon fluorescence imaging at isosbestic wavelength (810 nm). Lower, Myelin sheath wrapping optogenetically active axons visualized SCoRe (cyan). The axon diameters were estimated to by full-width-half-maximum of transverse intensity profile. (**b**) All axons in (a) consistently exhibited rise in intracellular calcium by optogenetic stimulation (shaded in magentha). (**c**) Averaged traces of activity-dependent change in diameter (ΔD) measured by SpeRe. The shaded cyan area indicates the standard error of the mean.

**Supplementary Table 1**

| Subcellular structure | Size | Refractive index |
| --- | --- | --- |
| Axon | 1 μm | 1.36 |
| Membrane | 5 nm | 1.50 |
| Cytoplasmic pocket | 200 nm | 1.35 |
| Cytosol (compact myelin) | 3 nm | 1.47 |
| Extracellular space (myelin) | 5 nm | 1.36 |
| Periaxonal space | 12 nm | 1.35 |
| # of myelin layers | 5 | - |

**Supplementary Table 1 | Optical parameter for numerical simulation.** The physical sizes and refractive indices of myelinated axons at the paranode used in the numerical simulation
